# A pathway for error-free non-homologous end joining of resected meiotic double-strand breaks

**DOI:** 10.1101/2020.06.17.157313

**Authors:** Talia Hatkevich, Danny E. Miller, Carolyn A. Turcotte, Margaret C. Miller, Jeff Sekelsky

## Abstract

Programmed DNA double-strand breaks (DSBs) made during meiosis are repaired by recombination with the homologous chromosome to generate, at selected sites, reciprocal crossovers that are critical for the proper separation of homologs in the first meiotic divisions. Backup repair processes can compensate when the normal meiotic recombination processes are non-functional. We describe a novel backup repair mechanism that occurs when the homologous chromosome is not available in *Drosophila melanogaster* meiosis. In the presence of a previously described mutation (*Mcm5*^*A7*^) that disrupts chromosome pairing, DSB repair is initiated by homologous recombination but is completed by non-homologous end joining (NHEJ). Remarkably, this process yields precise repair products. Our results provide support for a recombination intermediate recently discovered in mouse meiosis, in which an oligonucleotide bound to the Spo11 protein that catalyzes DSB formation remains bound after resection. We propose that this oligonucleotide functions as a primer for fill-in synthesis to allow scarless repair by NHEJ.

## INTRODUCTION

Crossovers promote the accurate segregation of homologous chromosomes during meiosis. The formation of crossovers begins with the introduction of double-strand breaks (DSBs) by the highly conserved endonuclease Spo11 and its associated proteins (reviewed in 1). Meiotic DSB repair differs from mitotic DSB repair in several important ways. First, DSB repair in mitotically proliferating cells can employ several repair strategies, including non-homologous end joining (NHEJ), various homology-directed pathways, and DNA polymerase theta-mediated end joining (reviewed in 2). In contrast, DSBs made during meiosis are, under normal circumstances, repaired exclusively by homologous recombination (HR). NHEJ begins with binding of the Ku heterodimer to DNA ends, thus it has been suggested that the bound Spo11 enzyme also blocks NHEJ by preventing binding of Ku (3,4). An early step in HR is 5’-to-3’ resection resulting in long, single-stranded 3’ overhangs that also block NHEJ (5-8).

A second key difference between mitotic and meiotic DSB repair is that the sister chromatid is typically used as a template for HR during mitotic repair but in meiosis the homologous chromosome is used. Bias for using the homolog is established by the presence of the meiotic chromosome axis and meiosis-specific recombination enzymes (9-11). The axis, made up of cohesins and meiosis-specific proteins, organizes the chromosome into an array of loops which later serve as the base of the synaptonemal complex (SC) that joins homologous chromosomes together (reviewed in 12).

Finally, mitotic HR is regulated to avoid generation of reciprocal crossovers but in meiosis reciprocal crossing over is essential and is actively promoted at selected DSB sites, the outcome of HR must, at selected sites (reviewed in 13). Complexes specific to meiotic recombination, including the ZMM proteins found in many eukaryotes (reviewed in 14) and the mei-MCM complex of flies (15), block non-crossover pathways and/or promote crossing over.

When meiotic recombination is disrupted by mutations in key genes, alternative repair pathways are activated to ensure that all DSBs get repaired. If HR is blocked at early steps, NHEJ and other pathways can intervene (5-8,16). If later steps in the meiotic crossover pathway are compromised, mitotic-like HR mechanisms take over (reviewed in 17).

Another interesting question is what happens when recombination pathways are intact but a homologous chromosome is not available to use as a template. In budding yeast and *Arabidopsis*, meiosis in haploids or when one chromosome is monosomic reveals that HR can be used as a repair template, though less efficiently (18-20).

We describe here a novel pathway for repair of meiotic DSBs when a homolog is not present. Hatkevich *et al*. (21) recently demonstrated that in *Drosophila melanogaster* females with the separation-of-function mutation *Mcm5*^*A7*^, chromosome pairing is established normally but is lost as the during assembly of the SC, resulting in heterosynapsis (SC between non-homologous chromosomes). Previous studies had found that DSBs appear and disappear with normal kinetics in *Mcm5*^*A7*^ mutants (22). Our studies show that DSB repair in these mutants is attempted first HR, but is then completed by NHEJ. Remarkably, whole-genome sequencing fails to find deletions predicted to be produced by NHEJ, indicating precise repair. We propose a model in which Spo11-bound oligonucleotides annealed to the ends of resected DSBs can function as primers for synthesis to fill in resected regions, allowing error-free repair by NHEJ. Recent studies provide evidence for a similar intermediate in mouse meiosis (23,24), suggesting that this may be a conserved mechanism for repair of meiotic DSBs when the homologous chromosome is unavailable.

## MATERIAL AND METHODS

### Genetic assays

Flies stocks were maintained on standard medium at 25°C. In this article, *Drosophila* nomenclature was generalized for readership. For transparency, genotypes, specific alleles, and associated figures are listed in the table below:

| Genotype in article | Specific genotype |
| --- | --- |
| WT | *y w<sup>1118</sup>* |
| *Mcm5<sup>A7</sup>* | *Mcm5<sup>A7</sup> / Df(3R)Exel7305* |
| *mei-P22* | *mei-P22<sup>103</sup>* |
| *mei-P22 Mcm5<sup>A7</sup>* | *mei-P22<sup>103</sup>, Mcm5<sup>A7</sup> / Df(3R)Exel7305* |
| *lig4* | *lig4<sup>57</sup>* |
| *lig4 mei-P22* | *lig4<sup>57</sup>; mei-P22<sup>103</sup>* |
| *lig4 Mcm5<sup>A7</sup>* | *lig4<sup>57</sup>; Mcm5<sup>A7</sup> / Df(3R)Exel7305* |
| *lig4 mei-P22 Mcm5<sup>A7</sup>* | *lig4<sup>57</sup>; mei-P22<sup>103</sup>, Mcm5<sup>A7</sup> / Df(3R)Exel7305* |
| *spnA* | *spnA<sup>093A</sup> / spnA<sup>057</sup>* |
| *Mcm5<sup>A7</sup> spnA* | *Mcm5<sup>A7</sup> / Df3R Exel7305, spnA<sup>093A</sup> / spnA<sup>057</sup>* |
| *mh* | *mh<sup>1</sup> / mh<sup>KG05829</sup>* |
| *mh Mcm5<sup>A7</sup>* | *mh<sup>1</sup> / mh<sup>KG05829</sup>; Mcm5<sup>A7</sup> / Df(3R)Exel7305* |
| *gkt Mcm5<sup>A7</sup>* | *P{nos::GAL4} / P{UAS::gkt RNAi}; Mcm5<sup>A7</sup> / Df(3R)Exel7305* |

*X* chromosome NDJ was evaluated by scoring progeny from virgin females of the desired genotype crossed with *y cv v f / T(1:Y)B*^*S*^ males. Viable exceptional *XXY* females have Bar-shaped eyes, and viable exceptional *XO* males have wild-type eye shape and express the *y cv v f* mutant phenotypes. Crossovers on chromosome *2L* were measured by crossing virgin *net dpp*^*d-ho*^ *dpy b pr cn /* + females of the desired genotype to *net dpp*^*ho*^ *dp b pr cn* males. Vials of flies were flipped after three days of mating. Resulting progeny were scored for all phenotypic markers.

### Dissection and immunofluorescence (IF) of whole mount germaria

In all immunofluorescent and genetic experiments, *Drosophila melanogaster* adult females 3-10 days old were used. In whole genome sequencing studies, individual male progeny were used. Ten three- to five-day old virgin females were fattened overnight with yeast paste in vials with ∼5 males of any genotype. Ovaries were dissected in fresh 1x PBS and incubated in fixative buffer for 20 minutes. Fixative buffer: 165 µL of fresh 1x PBS, 10 µL of N-P40, 600 µL of heptane, and 25 µL of 16% formaldehyde. After being washed 3 times in 1x PBS + 0.1% Tween-20 (PBST), ovaries were incubated for 1 hour in 1 mL PBST + 1% BSA (10 mL of PBST + 0.1 g BSA). Ovaries were incubated overnight in 500 µL primary antibody diluted in 1 mL PBST + 1% BSA at 4° C on a rocking nutator. Ovaries were then washed 3x in PBST and incubated in 500 µL secondary antibody diluted at 1:500 in PBST + 1% BSA for 2 hours under foil. Ovaries were mounted in 35 µL of ProLong Gold + DAPI on a microscope slide using fine-tip forceps to spread ovaries.

Antibodies for C(3)G (25) and g-H2Av (Rockland) were used. Images of whole mount germaria were taken Zeiss LSM880 confocal laser scanning microscope using 63x/0.65 NA oil immersion objective, with a 2x zoom using ZEN software. Images were saved as .czi files and processed using FIJI (26). DSBs were quantified as described below.

For quantification of g-H2Av foci, FIJI (26) was used to visualize images, and contrast and brightness were adjusted for optimal viewing. In each region of the germarium, individual g-H2Av foci were manually counted in nuclei expressing C(3)G. Data are represented as mean ± 95% CI, and the two tailed T-test or nested T-test was used to determine significance. Genotypes were blinded to the scorer.

### Whole genome sequencing, alignment, and SNP calling

*Mcm5*^*A7*^ was crossed into a *y cv v f* line with isogenized chromosomes *X* and *2* (*iso1*). *Df(3R)Exel7305* was crossed into a *w*^*1118*^ line with isogenized chromosomes *X* and *2*. These two lines were crossed to one another and resulting females (heterozygous for *iso1* and *w*^*1118*^ on *X* and *2* and *Mcm5*^*A7*^/*Df(3R)Exel7305* on *3*) were backcrossed with *iso1* males. Resulting male progeny were used for whole-genome sequencing.

DNA was isolated from individual male progeny using the Qiagen Blood and Tissue Kit and sheared using a Covaris S220 sonicator. KAPA HTP Library Prep kit (KAPA Biosystems, Cat. No. KK8234) was used to construct libraries with NEXTflex DNA barcodes (BiooScientific, Cat No. NOVA-514104). Libraries were pooled and run on an Illumina NextSeq 500 as 150 bp paired-end samples using the high-output mode. Real Time Analysis software version 2.4.11 and bcl2fastq2 v2.18 demultiplexed reads and generated FASTQ files. FASTQ files were aligned to version 6 of the *Drosophila melanogaster* reference genome (dm6) using bwa mem version 0.7.17-r1188 with default settings (27). SNPs were called using both Samtools (28) and GATK HaplotypeCaller (29). Depth of coverage analysis revealed two X0 males (mcm05-01 and mcm05-12).

### Detection of CO and NCO-GC events and calculation of expected events

CO and NCO-GC events were identified as in Miller *et al*. (30). Briefly, for each individual offspring sequenced, the VCF file was used to identify changes in polymorphisms from one parent to another. SNP density per chromosome arm was: 1/531 bp for the *X*, 1/255 bp for *2L*, 1/254 bp for *2R*, 1/249 bp for *3L*, and 1/318 bp for *3R*. Each candidate change was validated visually using IGV. A total of 28 individual males were sequenced to revealed 11 COs and 16 NCO-GCs. 11 NCO-GCs were validated using PCR and Sanger sequencing.

Expected CO and NCO-GC events were calculated based on wild-type values from Miller *et al*. (31) in which 541 CO and 294 NCO-GC events were observed in 196 wild-type meioses. Fisher’s exact test (two-tailed) was used to determine significance between observed and expected event frequencies (GraphPad QuickCalcs; http://www.graphpad.com/quickcalcs/).

### Identification of deletions and generation of simulated deletions

Multiple approaches were used to search for deletions. First, VCF files generated by Samtools and GATK HaplotypeCaller were searched for unique deletion polymorphisms. Candidate deletions were visually examined using IGV (32). Second, Pindel v0.2.5b9 (33) and Breakdancer v1.1_20100719 (34) were used to identify larger deletions and a custom script was used to parse the output and identify novel events. No novel deletions were identified using both approaches.

To validate our approach, simulations were done as in Miller (35). Briefly, 200 simulated genomes were created based on the dm6 reference genome, 100 with deletions of 1–20 bp, and 100 with deletions varying from 1–1,000 bp. SNPs were randomly placed approximately 1 every 500 bp in these synthetic genomes. For each genome a minimum of five and maximum of 25 deletions were made. Each deletion was randomly assigned a size, a position on *X, 2*, or *3*, and a one of the four haplotypes corresponding to the four meiotic products; one of these haplotypes was selected as the offspring. A second synthetic genome representing the male haplotype was also generated with no *X* but with a *Y* chromosome, with SNPs randomly placed approximately 1 every 500 bp. Using each synthetic genome as a reference, ART (36) was used to generate 100 bp paired-end reads at approximately 10x coverage with an average insert size of 400 bp for both the male and female synthetic genomes. FASTQ files were then combined into a single FASTQ and aligned to dm6 using bwa. These synthetic genomes were analyzed similar to the real ones, with SNPs called using both Samtools and GATK HaplotypeCaller, and Pindel and Breakdancer used to identify larger deletions. Analysis of vcf files from simulated genomes revealed that the approaches described above identify >95% of novel deletions of 1–20 bp and at least 65% of novel deletions 1–1,000 bp.

To calculate the number of deletions we expected to recover we used a conservative assumption of 15 DSBs per meiosis. Based on the whole-genome sequencing data, HR is decreased by ∼80% in *Mcm5*^*A7*^ mutants, suggesting that NHEJ repairs at least 12 DSBs per meiosis; each oocyte receives one of four chromatids, resulting in an average of three deletions per progeny. Therefore, we expect 84 deletions in our sample size of 28 meioses. Based on the simulated genomes described above, we should detect 71 of these if they are small, 48 if they are large.

Normal and exceptional (those resulting from a nondisjunction event) progeny were counted. To adjust for inviable exceptional males and females, viable exceptional class was multiplied by 2. % *X*-NDJ = 100* ([2*viable exceptional females] + [2*viable exceptional males])/total progeny. Statistical comparisons were performed as in Zeng et al (37).

Genetic distances are expressed in centiMorgans (cM), calculated by 100 * (*R* / *n*), where *R* is the number of recombinant progeny in a given interval (including single, double, and triple crossovers), and *n* is the total number of progeny scored. 95% confidence intervals were calculated from variance, as in Stevens {Stevens, 1936 #5180}]. Molecular distances (in Mb) are from the positions of genetic markers on the *Drosophila melanogaster* reference genome, release 6.12 (38). Crossover density (or frequency), as calculated by cM/Mb, exclude transposable elements (39).

## RESULTS

### Reduction of crossovers and loss of crossover patterning in Mcm5A7 mutants

*Mcm5*^*A7*^ is a missense, separation-of-function mutation recovered in a screen for mutants causing high levels of meiotic nondisjunction (22). The high level of nondisjunction observed in *Mcm5*^*A7*^ mutants was attributed to a 90% reduction in meiotic crossovers on the *X* chromosome. The decrease in crossing-over was non-uniform with the interval spanning the centromere being decreased by only 80%. We extended this analysis to an autosome by scoring crossovers along *2L* and observe an overall decrease of about 75%. However, the nonuniformity is more pronounced as one interval (*b* to *pr*) exhibits 150% of crossovers compared to wild-type flies (Figure 1A, Table S1). Notably, in wild-type flies this interval accounts for about 10% of the crossovers on *2L*, while in *Mcm5*^*A7*^ mutants more than half of all crossovers are in this region.

**Figure 1.**
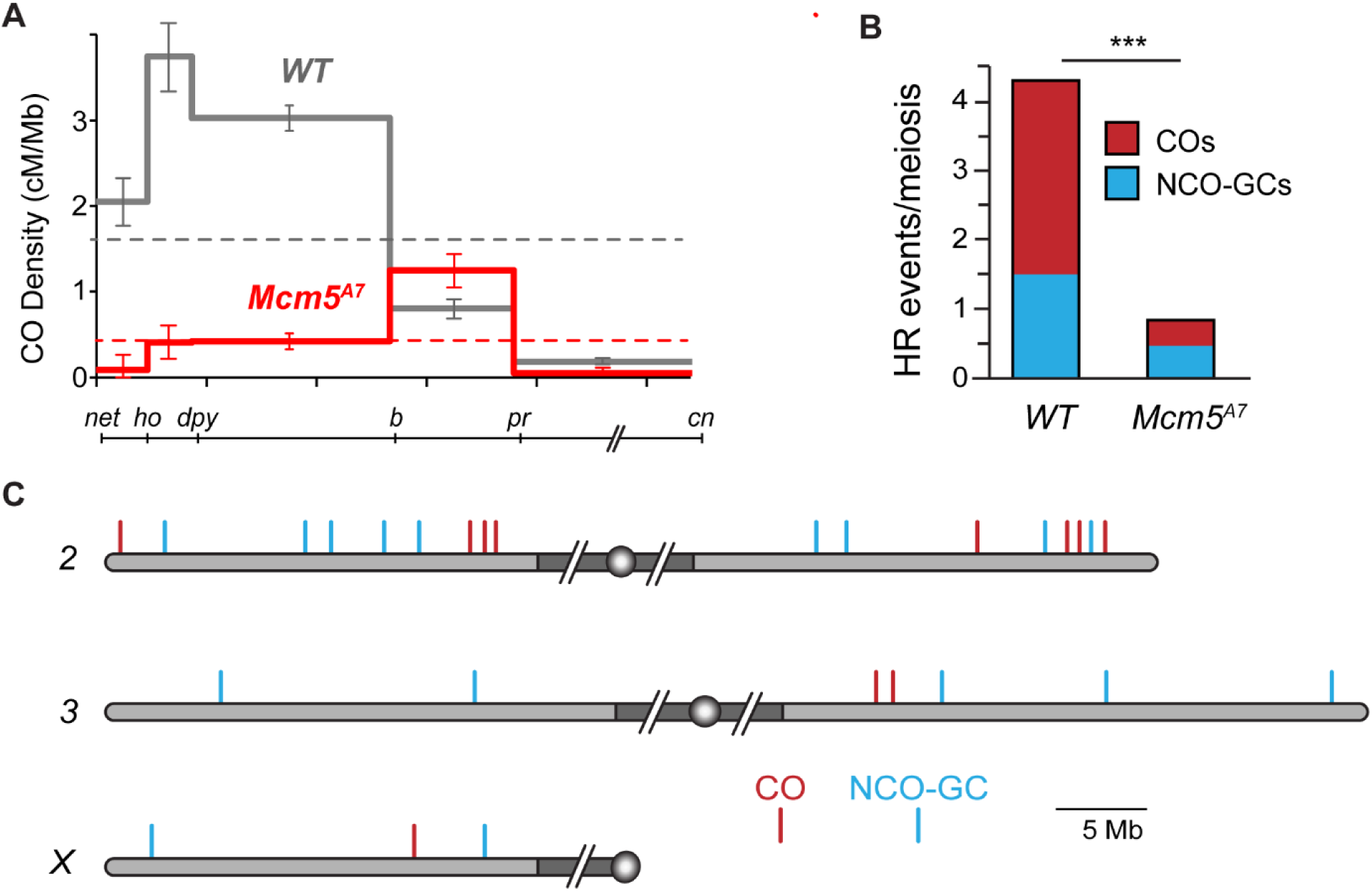
Homologous recombination is severely reduced in *Mcm5*^*A7*^ mutants. (**A**) Crossover density (cM/Mb) across *2L* in wild-type females and *Mcm5*^*A7*^ mutants (*n* = 2070). Dashed lines represent mean crossover density across the region assayed. Error bars indicate 95% confidence intervals. The line below the graph shows the markers used along *2L* and *2R* (slashes represent the centromere and pericentric heterochromatin, which are not included). (**B**) Number of crossovers (COs) and non-crossover gene conversion events (NCO-GCs) per meiosis in wild type (*n* = 196 meioses, dataset from Miller *et al*. (31) and *Mcm5*^*A7*^ mutants (*n* = 28 meioses). \*\*\**p* < 0.0001, T-test, two tailed. **(C)** COs and NCO-GCs identified by WGS of individual males from *Mcm5*^*A7*^ females. Gray shaded area represents euchromatic sequence, while dark gray shaded area represents the pericentric heterochromatin (not to scale), circles represent centromeres. *2L* is vertically aligned with the graph in panel (**A**).

Crossover distribution is a product of meiotic spatial patterning phenomena, especially crossover interference. Interference is observed as a decrease in the frequency of double crossovers from the number expected if crossovers are independent of one another (reviewed in 40). Because crossover distribution is perturbed in the *Mcm5*^*A7*^ mutant, we asked whether interference is affected. Due to the strong reduction in crossovers in the *Mcm5*^*A7*^ mutant, there is only one pair of adjacent intervals in our dataset with enough crossovers to estimate interference: *dp* to *b* and *b* to *pr* (Table S1). In wild-type flies, the incidence of double crossovers in these two intervals is less than half the number expected if they are independent (39). In *Mcm5*^*A7*^ mutants, however, six double crossovers were expected and seven were observed, consistent with a partial or complete loss of interference.

The SC has been suggested to be important for achieving interference (40,41). The effect on interference may be explained by the presence of heterosynapsis in *Mcm5*^*A7*^ mutants (21); we hypothesize that discontinuities in the SC (*i*.*e*., heterosynapsis to homosynapsis) impede interference. The higher incidence of crossovers in the *b* to *pr* interval is intriguing. In *Mcm5*^*A7*^ mutants, chromosome pairing appears to be normal in late leptotene is disrupted as SC assembles. The higher frequency of crossovers in the *b* to *pr* region might result from greater stability of pairing in this region.

### Homologous recombination is severely reduced in Mcm5A7 mutants

Because the number of meiotic DSBs in *Mcm5*^*A7*^ mutants is normal (22, see below), the reduction in crossovers in *Mcm5*^*A7*^ mutants likely reflects either a direct role for Mcm5 in the crossover pathway or a reduced ability to use the homologous chromosome for HR. In wild-type females, about a third to a quarter of DSBs are repaired as crossovers, with the remainder becoming non-crossovers; only crossovers would be reduced if these mutants have a crossover-specific defect, but both crossovers and noncrossovers would be reduced if the defect is in accessing the homologous chromosome. We used whole-genome sequencing to measure both crossovers and non-crossovers in individual offspring from *Mcm5*^*A7*^ females. Since only one of four chromatids goes into the oocyte, only half of the crossover and one quarter of the non-crossovers will be detected. Only those non-crossovers that are accompanied by a gene conversion tract that spans at least one polymorphism (non-crossover gene conversions; NCO-GCs) will be detected.

We whole-genome sequenced 28 individual progeny from *Mcm5*^*A7*^ females, identified crossover and NCO-GC events, and compared this data to a previously published dataset of 196 individual progeny from wild-type females (31). Genome-wide, we find an 86% reduction in crossovers (78 expected vs 11 observed, *p*<0.001) and a 67% reduction in NCO-GCs (42 expected vs 14 observed, *p*<0.001, two-tailed T-test) (Figure 1B). A reduction in both crossovers and NCO-GCs is consistent with a general inability to complete interhomolog HR in *Mcm5*^*A7*^ mutants.

Given the results with traditional crossover mapping described above, we also asked whether crossovers and NCO-GCs cluster in the genome. On *2L*, three of the four crossovers observed are located within a 2 Mb span between *b* and *pr*, the region in which genetic experiments also revealed an elevated rate of exchange (Figure 1C). Similarly, on *2R*, three of the four crossovers are grouped within a 2 Mb region, and the two COs on *3R* occur within 0.4 Mb of one another (no COs were recovered on *3L* and only a single CO was recovered on the *X*). In contrast, there is no obvious clustering of NCO-GCs, which appear to be evenly distributed throughout the genome (Figure 1C). These findings suggest that CO and NCO formation may be differentially controlled in *Mcm5*^*A7*^ mutants.

### Most double-strand breaks are repaired with normal kinetics in *Mcm5*^*A7*^ mutants

Lake *et al*. (22) found that meiotic DSBs, as marked by γ-H2Av foci, are created and repaired with normal kinetics in *Mcm5*^*A7*^ mutants. To extend this result, we counted γ-H2Av foci during DSB formation and repair. Foci first appear in Region 2A of the germarium, then decrease in number as the pro-oocyte progresses through Region 2B and into Region 3, with few or no foci remaining in Region 3, by which time breaks are thought to be repaired (Figure 2A).

**Figure 2.**
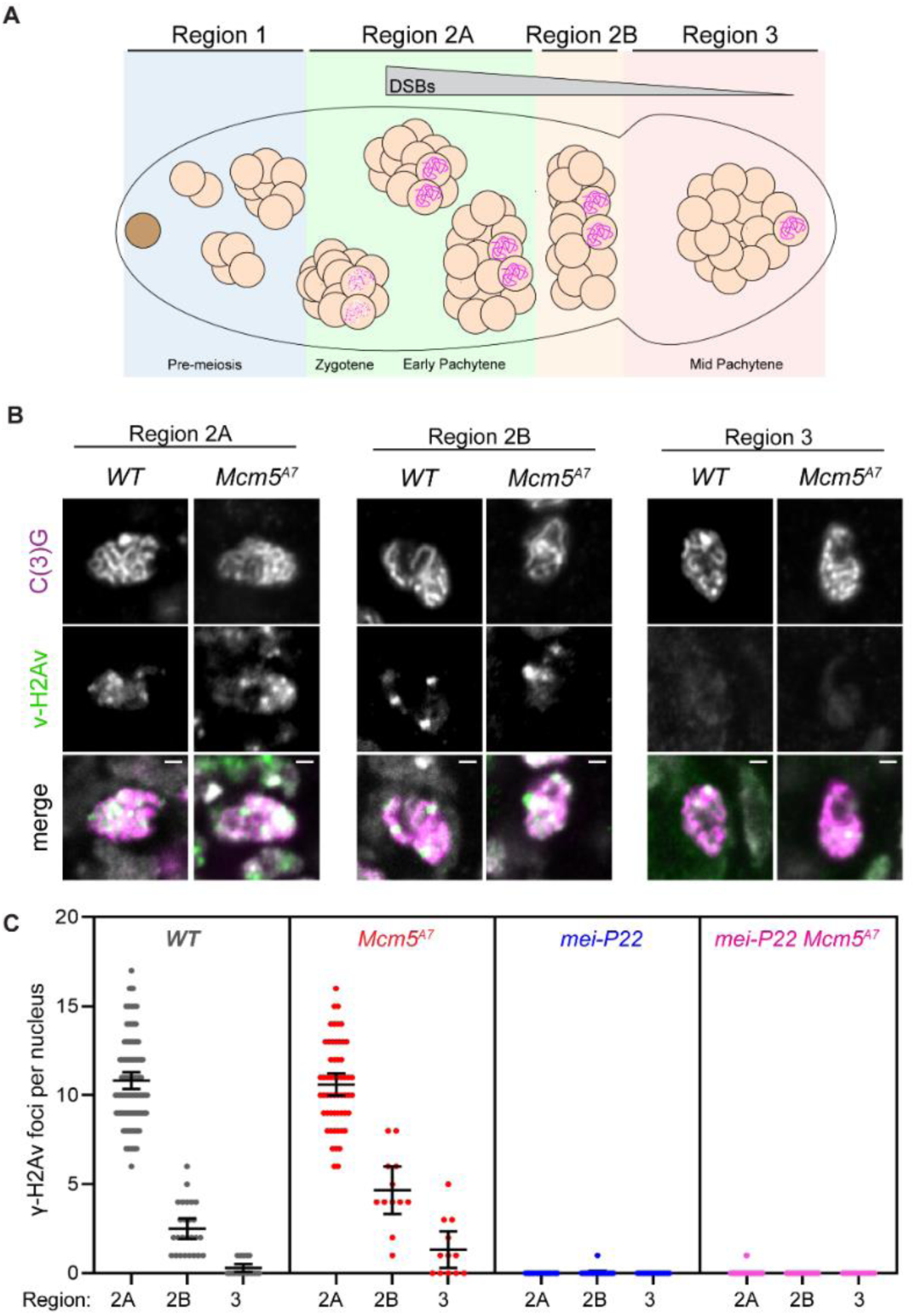
Meiotic double-strand breaks are repaired with normal kinetics in *Mcm5*^*A7*^. (**A**) Schematic of the *Drosophila* germarium. The germline stem cell (brown), cystoblast (not shown), and developing 2-, 4-, and 8-cell cystocytes (cc) reside in the pre-meiotic mitotic region of the germarium, Region 1. Meiotic onset occurs within the most anterior 16-cc in Region 2A, which is cytologically defined as zygotene. In early pachytene (Region 2A), formation of the synaptonemal complex (SC, pink scribble) is complete and meiotic double-strand break (DSB) formation begins. Meiotic DSBs are repaired as the 16-cc progresses through early pachytene (Region 2B) and into mid-pachytene (Region 3). Gray puncta in merged images are DAPI-dense chromocenters. (**B**) Representative images of g-H2Av foci (green) in *WT* and *Mcm5*^*A7*^ meiotic nuclei (as defined by the present of the SC component, C(3)G, pink) within Regions 2A, 2B, and 3. Merge images include DAPI in gray. Image brightness and contrast have been adjusted for clarity. (**C**) Quantification of g-H2Av foci. Each dot represents one nucleus; bars show mean and 95% confidence intervals. *n* = (across the three regions) for *WT*: 99, 25, 18; for *Mcm5*^*A7*^: 48, 10, 10; for *mei-P22*: 60, 19, 11; for *mei-P22 Mcm5*^*A7*^: 68, 19, 11.

*Mcm5*^*A7*^ mutants have a similar number of γ-H2Av foci in Region 2A as wild type (Figure 2B, 2C). Given the canonical function of Mcm5 in DNA replication (42), it seemed possible that some DSBs in *Mcm5*^*A7*^ mutants result from problems during pre-meiotic S phase. To test this, we used a *mei-P22* mutation to eliminate meiotic DSBs. γ-H2Av foci are nearly eliminated in both *mei-P22* single mutants and *mei-P22 Mcm5*^*A7*^ double mutants (Figure 2B, 2C), indicating that breaks within *Mcm5*^*A7*^ mutants are meiotically-induced.

The number of γ-H2Av foci in Regions 2B and 3 is slightly higher in *Mcm5*^*A7*^ mutants than in wild-type females (*p* = 0.0006 and *p* = 0.0112, T-test, two-tailed). The mean of 10.6 foci in Region 2A likely underestimates the total number of DSBs per meiotic cell due to asynchrony in formation and repair. Estimates range from 11 to 24 based on γ-H2Av foci and whole-genome sequencing (31,43). Thus, we estimate that 90-95% of DSBs made in *Mcm5*^*A7*^ mutants are repaired by Region 3. Since interhomolog HR is reduced by >80% (Figure 1), these DSBs must be repaired in other ways.

### Repair of most DSBs in Mcm5^A7^ mutants requires both Rad51 and DNA ligase IV

Previous studies have shown that *Mcm5*^*A7*^ mutants do not exhibit an increase in inter-sister crossing-over (6,21) suggesting that a strong barrier to use of the sister as an HR repair template is present in regions of heterosynapsis as in regions of homosynapsis. Together, our data suggest that DSBs made in regions of heterosynapsis are repaired by one or more non-HR pathways. One possibility is NHEJ. NHEJ is normally prevented during meiotic DSB repair but can function when there are defects in HR (5-8,16,44). To determine whether NHEJ is responsible for meiotic break repair in *Mcm5*^*A7*^ mutants, we asked whether repair is dependent on DNA ligase IV (Lig4), an enzyme both specific to and essential for NHEJ (45). In *lig4* single mutants, γ-H2Av foci in Regions 2A, 2B, and 3 are similar to those in wildtype (Figure 3A, panel 1, *p* = 0.8048 versus wildtype, nested t-test, two-tailed). As in wild-type and *Mcm5*^*A7*^ mutants, most γ-H2Av foci in *lig4* mutants are dependent on Mei-P22 (Figure 3A, panel 2). These results confirm previous observations that Lig4 is not involved in normal meiotic break repair in *Drosophila* (6). In *lig4 Mcm5*^*A7*^ double mutants, the number of γ-H2Av foci in Region 2A is similar to that seen in wild-type and single mutants, but unlike *Mcm5*^*A7*^ single mutants, most foci persist through Region 2B and into Region 3 (Figure 3A, Panel 3). Most of these foci are dependent upon Mei-P22 (Figure 3A, Panel 4). Thus, the timely repair of meiotic DSBs in *Mcm5*^*A7*^ mutants requires Lig4.

**Figure 3.**
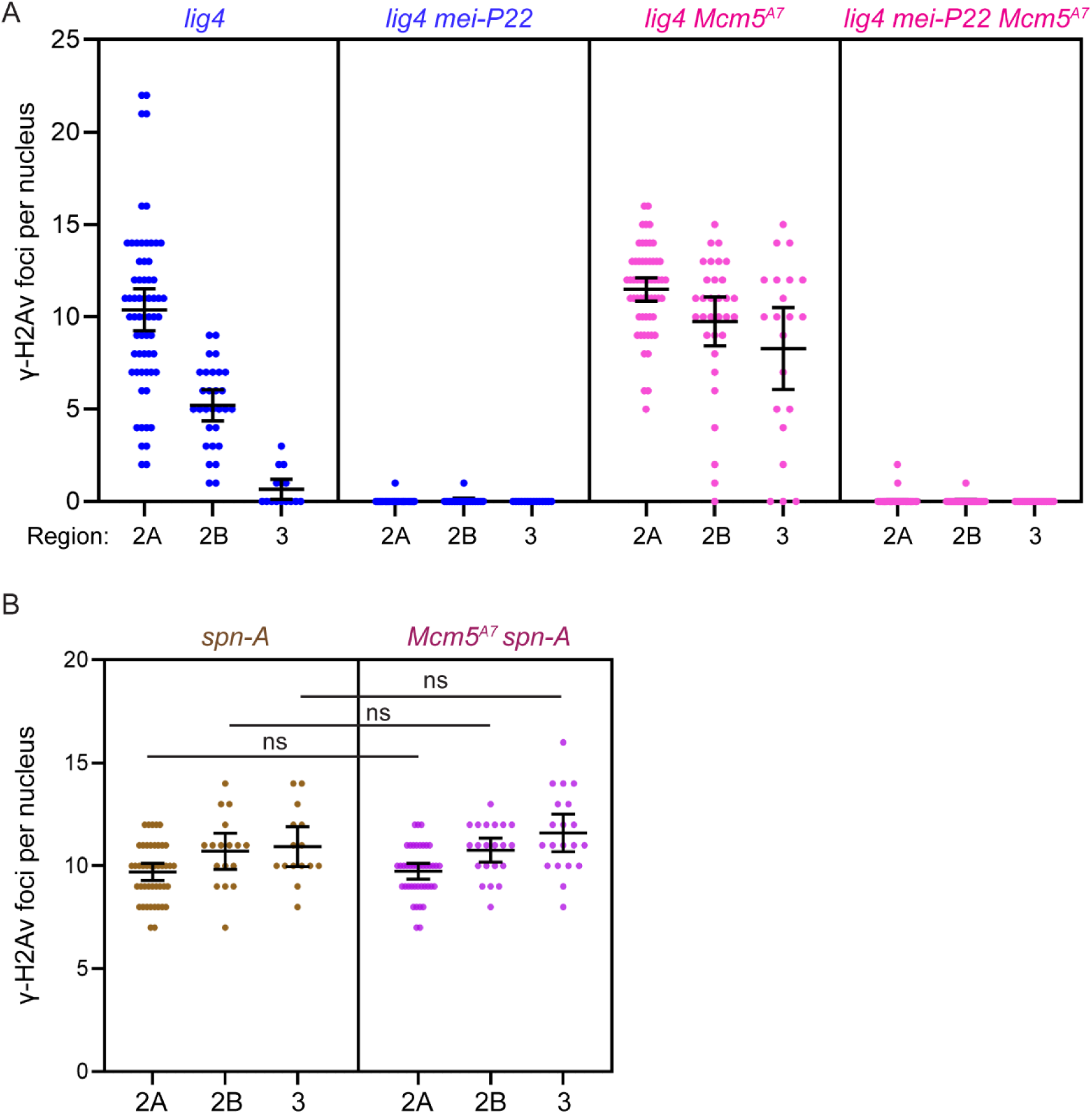
Meiotic DSB repair in *Mcm5*^*A7*^ mutants requires the NHEJ protein Lig4 and the strand exchange protein Spn-A. (**A**) and (**B**) Quantification of g-H2Av foci. Each dot represents one nucleus; bars show mean and 95% confidence intervals. *n* = (across the three regions) for *lig4*: 63, 29, 15; for *lig4 mei-P22*: 46, 17, 14; for *lig4 Mcm5*^*A7*^: 57, 32, 21; for *lig4 mei-P22 Mcm5*^*A7*^: 74, 24, 17; for *spn-A*: 44, 17, 15; for *Mcm5*^*A7*^ *spn-A*: 42, 21, 20.

The most straightfoward interpretation of our γ-H2Av focus data is that there can be both homosynapsis and heterosynapsis within a single nucleus, and that DSBs in regions of homosynapsis are repaired by HR, whereas DSBs in regions of heterosynapsis are repaired by NHEJ. An important question is whether HR is initially attempted at all DSBs, with NHEJ as a backup. To address this question, we determined what proportion of DSBs in *Mcm5*^*A7*^ mutants are dependent on Rad51 (in *Drosophila*, this is encoded by the *spn-A* gene; for simplicity, we refer to the Spn-A protein as Rad51). Meiotic DSBs are not repaired in *spn-A* mutants, resulting in a modest accumulation of foci as development proceeds (Figure 3B). Surprisingly, *Mcm5*^*A7*^ *spn-A* double mutants have the same number of γ-H2Av foci in all regions as *spn-A* single mutants (Figure 3B). This implies that all breaks, regardless of whether located in regions of heterosynapsis or homosynapsis, are initially processed by HR.

### NHEJ-dependent DSB repair in *Mcm5*^*A7*^ mutants is precise

There are several steps in the meiotic recombination pathway at which NHEJ might be enabled. Meiotic DSBs are formed from when Spo11 (*mei-W68* in *Drosophila*; for simplicity, we refer to the Mei-W68 protein as Spo11) cleaves with 2-bp 5’ overhanging ends to which the enzyme remains covalently bound (46) (Figure 4A). It is thought that Spo11 blocks binding of Ku, the first step in NHEJ (3,4). Clipping of the 2-nt overhangs by an endonuclease would remove this block but subsequent repair by NHEJ would then result in a deletion of at least two base pairs (Figure 4A). In most models, Spo11-bound oligonucleotides are released during resection (Figure 4B). NHEJ cannot function on long ssDNA overhangs. For example, the 17-nt overhangs generated during excision of *P* transposable elements appear to be refractory to NHEJ (47). These overhangs might also be clipped by and endonuclease (or possibly degraded by an exonuclease); NHEJ would then result in larger deletions, perhaps in the 50-100 bp range (Figure 4B). Finally, for NHEJ to function after long resection the long ssDNA regions would need to be removed, and this would generate large deletions (Figure 4C). We do not know the length of resection in *Drosophila*, but average gene conversion tract length of 350-450 bp (31,48,49) suggest a lower limit in this range, or perhaps twice this size is this reflects resection on only one side of the break and resection is symmetric. The requirement for Rad51 to repair all DSBs suggests that resection does indeed occur.

**Figure 4.**
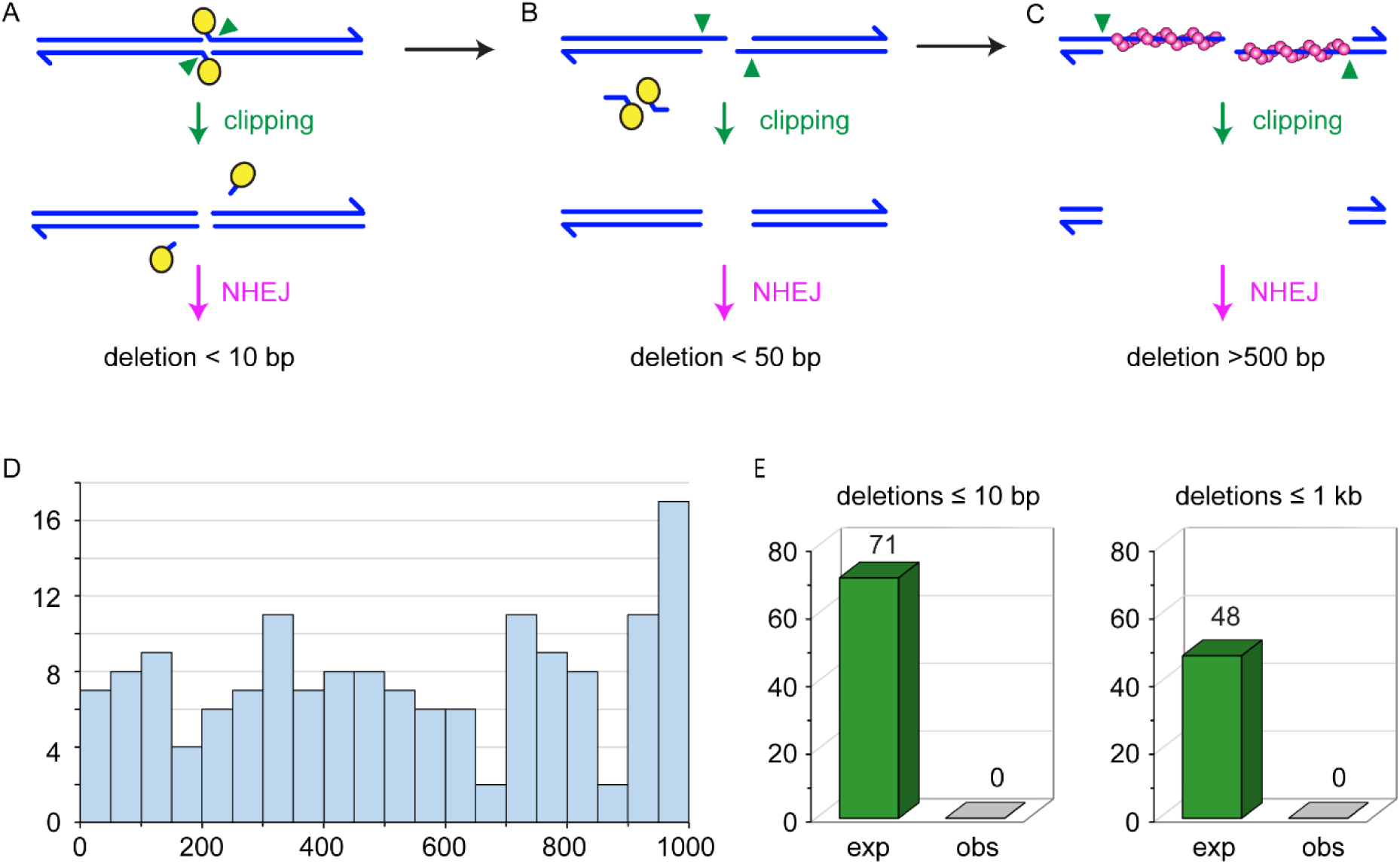
Expected and observed deletions. (**A-C**) Running across the top are structures from canonical models of meiotic recombination. Blues lines represent the two strands of a DNA molecule that is cleaved by Spo11 (yellow ovals). Spo11 remains covalently bound to the two-nucleotide overhanging 5’ ends. These overhangs might be clipped (green arrowheads) to produce ends on which NHJE can act. The repaired chromatid would have a deletion of at least two base pairs, but likely less than ten base pairs. (**B**) During normal resection, Spo11 is released bound to 16-24 nucleotide strands. If this occurs through initial nicks, overhangs of 14-22 nucleotides would be produced, with two base pairs lost. It may be possible that such overhangs could be joined by NHEJ, possibly with additional deletion, but it is likely that clipping would be required to produce a NHEJ substrate; this would result in larger deletions. (**C**) After long resection, Rad51 (magenta spheres) is loaded onto the ends. Given the requirement for Rad51/Spn-A in our experiments, this stage is likely reached at all DSBs. If homology search is unsuccessful, this long overhangs could be clipped to produce a substrate amenable to NHEJ, but the result would be a deletion that, based on gene conversion tract lengths, is would likely be more than 500 base pairs. (**D**) Histogram of sizes of deletions recovered. Each bin spans 50 bp. (**E**) Expected and observed deletions. Based on detection efficiency in simulations, we expected to detect 71 deletions in our dataset if deletions are less than ten base pairs, and 48 deletions if they are less than 1000 base pairs. We detected zero deletions of either class.

We used the methods described in Miller (35) to search for *de novo* deletions in the *Mcm5*^*A7*^ whole-genome sequence dataset described above. In simulated sequencing of artificial genomes with small deletions (1-10 bp), 85% were detected; in similar simulations with large deletions (1-1000 bp), 57% were detected, with no apparent bias in size within this range (Figure 4D). These methods successfully identified 20 *de novo* deletions of various sizes in sequences from progeny of mutants with defects in a synaptonemal complex protein (35). The 28 progeny we sequenced would be expected to harbor approximately 84 sites repaired by NHEJ (see Methods for calculation details). Based on simulations, we would expect to detect 71 of these if all of them are small, or 48 if they are under one kb (Figure 4E). We did not detect any deletions in our dataset. We also looked for translocations or inversions that might result from joining of misappropriate ends, using methods that have been used successfully to map rearrangements in balancer chromosomes (50,51); we did not detect any such rearrangements. Taken together, we conclude that NHEJ is able to repair ends precisely in *Mcm5*^*A7*^ mutants.

## DISCUSSION

Previous studies have found that loss of proteins required to catalyze the resection needed for HR can lead to repair of meiotic DSBs by NHEJ (5-8). Here, we have shown that most DSBs made in *Drosophila Mcm5*^*A7*^ mutants require both Rad51 and Lig4, indicating that NHEJ can be used to repair DSBs even after resection, presumably at sites where the homologous chromosome is not available as a template. Although NHEJ is generally characterized as error-prone, in this situation it is precise, with no detectable deletion.

We propose a model for precise NHEJ post-resection based on the recent discovery that Spo11-oligonucleotides remain annealed to DSB ends after resection in mouse meiosis (23,24) (Figure 5). In our model, the Spo11-oligonucleotide remains annealed during the homology search (Figure 5a-c). If a homologous template is found, strand exchange produces a D-loop (Figure 5d), then the Spo11-oligonucleotide are released by dissociation, cleavage invading strand to remove the double-stranded region, or enzymatic reversal of the tyrosyl phosphodiesterase bond. This allows repair synthesis to proceed (Figure 5e). We hypothesize that if homology cannot be found, such as in regions of heterosynapsis in *Mcm5*^*A7*^ mutants, the Spo11-oligonucleotides are used as primers for fill-in synthesis (Figure 5f). The Spo11 is then enzymatically removed and NHEJ proceeds through annealing of the 2-nt overhangs (Figure 5g) to restore the original sequence (Figure 5h).

**Figure 5.**
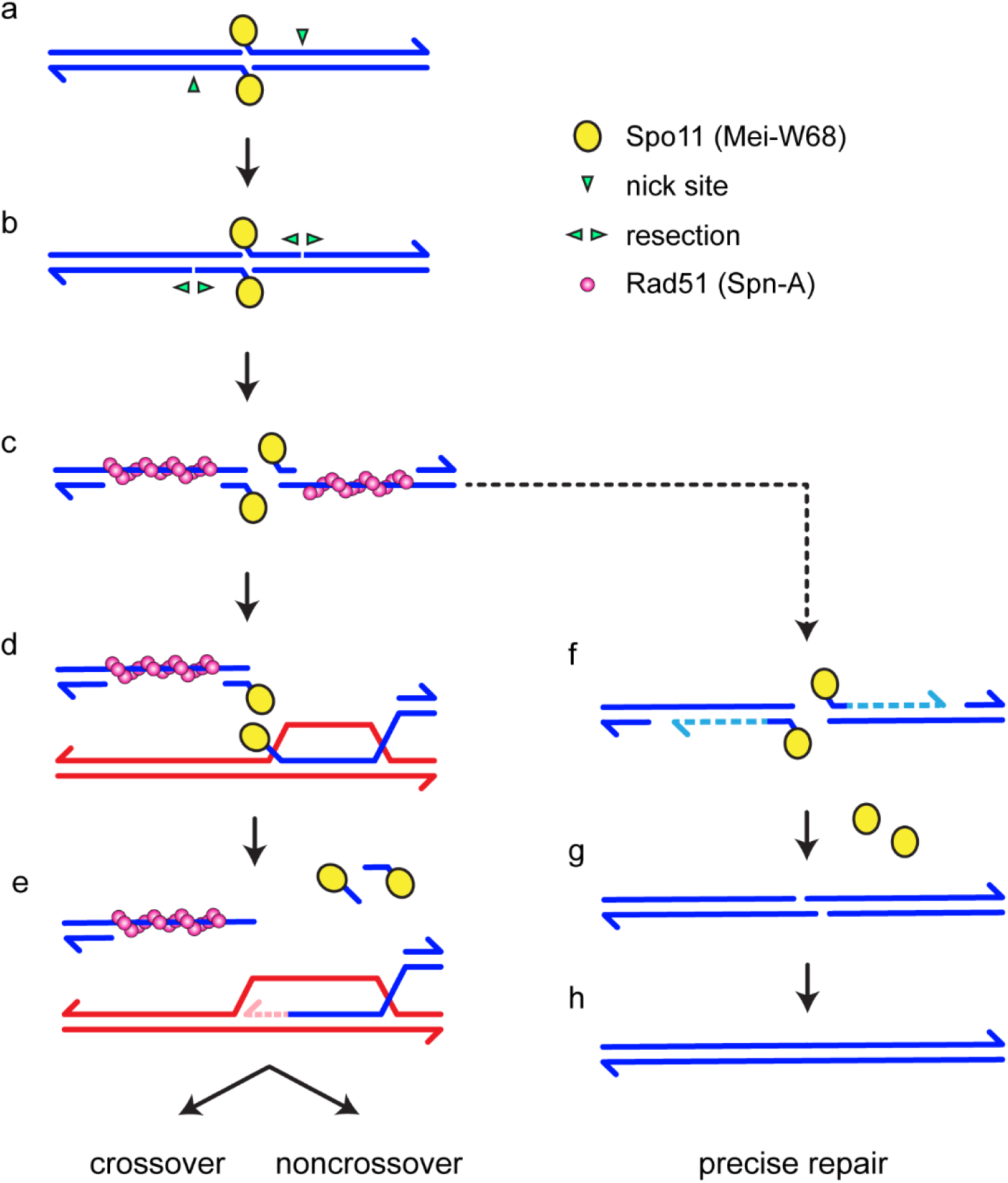
Models for Rad51-, Lig4-dependent meiotic DSB repair. See Figure S1 for additional models. In the canonical model (left), DSB formation involves concerted nicking by two Spo11 proteins (yellow circles), which remain covalently bound to 2-nt 5’ overhangs through a tyrosyl phosphodiesterase bond (a). Resection is initiated by nicks (green arrowheads) 3’ of the DSB, then (b) extended bidirectionally (3). Rad51 and associated proteins (and in many species, Dmc1) assembles on the single-stranded DNA exposed by resection (c). Recent observations suggest that a short Spo11-oligonucleotide remains annealed at the 5’ end of each strand (23,24). A homology search and strand exchange produces a D-loop (d). Removal of the Spo11-oligonucleotides, either by cleavage or dissociation, allows repair synthesis to proceed (e). Subsequent steps (not shown) produce crossover and non-crossovers. We hypothesize that if a homologous template cannot be found, such as in regions of heterosynapsis in *Mcm5*^*A7*^ mutants, the Spo11-oligonucleotides function as primers for fill-in synthesis (f). NHEJ can then complete repair through annealing of the 2-nt overhangs (g), resulting in precise repair (h).

A key unknown in our model is how Spo11 is removed. Likely candidates include tyrosyl-phosphodiesterases (TDPs), which directly reverse protein tyrosyl-DNA complexes generated by topoisomerases (reviewed in 52), and proteins with an SprT metalloprotease domain that can remove DNA-protein crosslinks (53-55). The only TDP in *Drosophila* is Tdp1, encoded by the gene *glaikit* (*glk*) (56,57). Because *glk* is an essential gene, we generated flies with germline expression of a transgene for RNAi knockdown of *glk* in *Mcm5*^*A7*^ mutants. We did not detect persistence of γ-H2Av foci in these females (Figure S2).

*Drosophila* has orthologs of the SprT domain proteins SPRTN and Germ Cell Nuclear Acidic Peptidase (GCNA). The SPRTN ortholog is encoded by *maternal haploid* (*mh*) (58). We constructed *mh Mcm5*^*A7*^ double mutants but did not detect persistence of γ-H2Av foci (Figure S2). The GNCA ortholog is difficult to test because mutants have Spo11-independent DSBs that apparently stem from defects in pre-meiotic S phase, as well as germline developmental defects in the ovary (59).

We do not know whether loss of the ability to remove Spo11 by a TDP or protease would lead to persistence of γ-H2Av foci in *Mcm5*^*A7*^ mutants. Blocking this step might allow cleavage of the 2-nt overhang, as Figure 4A. If this is the case, then foci might still go away with normal kinetics due to NHEJ repair; however, there would be a small deletion at every site that is unable to complete HR. This might provide a way to map DSBs in *Drosophila*.

In summary, our experiments reveal the existence of a novel backup mechanism to repair meiotic DSBs when an HR template cannot be found. This involves repair by NHEJ that, even though resection apparently occurs, is precise. We propose that this is possible because of retention of Spo11-bound oligonucleotides at the ends of the DSBs. Our results, coupled with the discovery that Spo11-oligos remain anneal in mouse meiosis (23,24), suggests that this backup pathway may be conserved in animals.

## AVAILABILITY

Illumina reads used in this project have been deposited at the National Center for Biotechnology Information (https://www.ncbi.nlm.nih.gov/) under project PRJNA623030. Code used in this project are available at: https://github.com/danrdanny/mcm5-A7.

## ACCESSION NUMBERS

Illumina reads used in this project have been deposited at the National Center for Biotechnology Information (https://www.ncbi.nlm.nih.gov/) under project PRJNA623030.

## SUPPLEMENTARY DATA

Supplementary Data include Figures S1 and S2.

## ACKNOWLEDGEMENT

We thank Scott Keeney for insightful discussion, the Sekelsky lab for helpful comments on the manuscript, and the Molecular Biology core at the Stowers Institute for Medical Research for expert help with DNA sequencing.

