## Supplementary material for "A pathway for error-free non-homologous end joining of resected meiotic double-strand breaks": Table S1, Figures S1, S2

**This PDF file includes:**

Table S1

Figures S1 and S2

**Table S1. Meiotic crossovers on chromosome *2L*.**

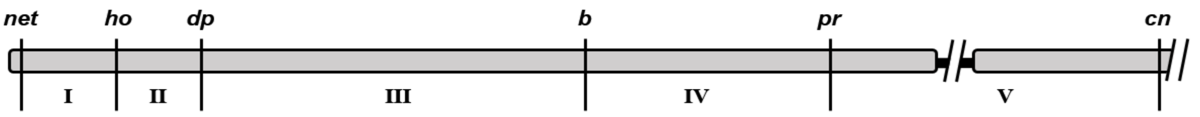


| Progeny Class | | Maternal Genotype | |
|---|---|---|---|
| | | *wild type* | *Mcm5$^{A7}$* |
| Parental | | 2376 | 1829 |
| Single crossover (interval) | I | 176 | 4 |
| | II | 290 | 14 |
| | III | 1099 | 16 |
| | IV | 154 | 135 |
| | V | 39 | 6 |
| Double crossover (intervals) | I / II | 1 | 0 |
| | I / III | 11 | 0 |
| | I / IV | 10 | 0 |
| | I / V | 2 | 0 |
| | II / III | 6 | 1 |
| | II / IV | 7 | 2 |
| | II / V | 13 | 0 |
| | III / IV | 19 | 7 |
| | III / V | 17 | 1 |
| | IV / V | 2 | 2 |
| | Totals: | 1903 | 2070 |

**Table S1. Meiotic crossovers on chromosome *2L*.** Each row lists the number of total progeny from parental, single crossover, and double crossover classes for wild-type and *Mcm5A7* mutant females. Wild-type data are from Hatkevich *et al.* (2017).

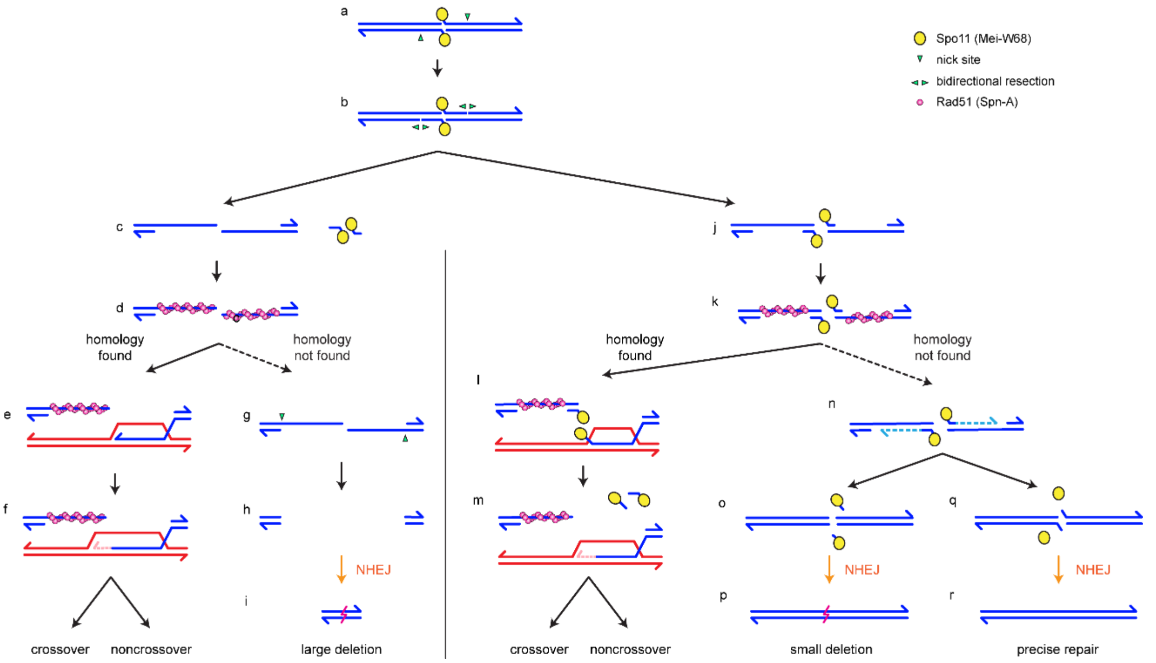


**Fig. S1.** Models for post-resection repair of meiotic DSBs by NHEJ. DSB formation by Spo11 (a) is followed by nicking (b). In most models (left), bidirectional resection that leads to release of Spo11-bound oligonucleotides (c). Rad51 and related proteins are loaded onto the ssDNA (d) and a homology search is done. After a successful homology search, strand exchange generates a D-loop (e) that is extended by synthesis (f). Further steps (not shown) generate crossover or noncrossover products. If a homologous recombination partner is not found (dashed arrow), the overhangs must be nicked (g, green arrowhead), leading to a large gap (h). The ends can now be joined by canonical NHEJ, resulting in a deletion likely to span 100s of base pairs (i). Models on the right are based on recent observations that suggest that Spo11-oligonucleotides remain bound to the 3' ends after resection (j). The Rad51 filament then spans a region of ssDNA gap (k). Successful strand exchange does not involve the extreme 3' end (l), but release of the Spo11-oligonucleotides by dissociation or cleavage would allow repair synthesis (m). As in the standard model, further steps lead to crossover or noncrossover products. If the homology search fails (dashed line), the Spo11-bound oligonucleotide could serve as a primer for gap-filling synthesis (n). Removal of Spo11 can be accomplished by clipping the 2-nt overhangs (o). In this case, NHEJ will result in a small deletion (p). Alternatively, Spo11 can be removed by reversal of the tyrosyl phosphodiesterase bond (q). Repair by NHEJ using the complementary 2-nt overhangs would restore the original sequence precisely (r). Our failure to detect either small or large deletions under conditions where NHJE is completing repair (*i.e.*, in the *Mcm5*$^{A7}$ mutant) , is most consistent with this last model.

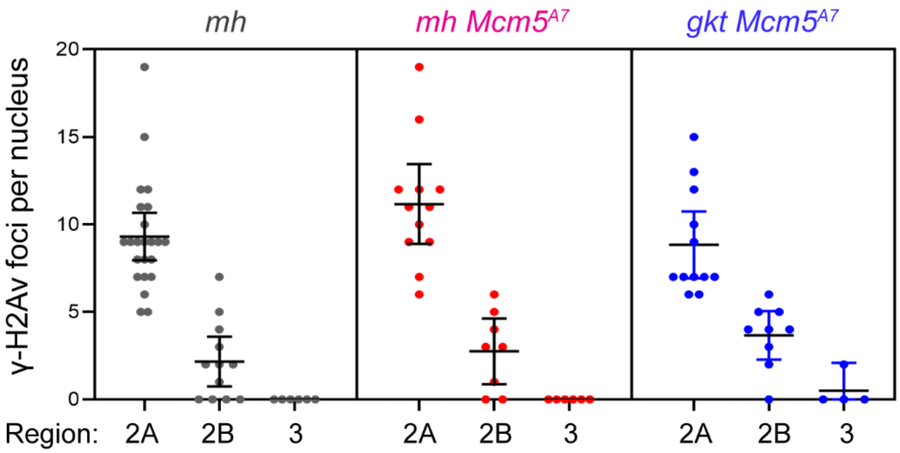


**Fig. S2.** Meiotic DSB repair in *Mcm5*$^{A7}$ mutants with potential Spo11 nucleases compromised. Each dot is the quantification of γ-H2Av foci in one nucleus; bars show mean and 95% confidence intervals. *n* = (across the three regions) for *mh*: 23, 19, 6; for *mh Mcm5*$^{A7}$: 12, 8, 6; for *gkt Mcm5*$^{A7}$: 12, 10, 4. The genotype listed *mh* is *mh*; [*Mcm5*$^{A7}$ or *Df(3R)Exel7305*] / *TM6B*. The *gkt* in the genotype above is a combination of RNAi line and *nos::GAL4*, which expresses before meiosis begins and throughout pachytene (in the absence of an antibody we had no way to quantify knockdown of GKT protein).
